# Mining transcriptomic data to identify *Saccharomyces cerevisiae* signatures related to improved and repressed ethanol production under fermentation

**DOI:** 10.1101/2021.10.21.465282

**Authors:** Sima Sazegari, Ali Niazi, Zahra Zinati, Mohammad Hadi Eskandari

## Abstract

*Saccharomyces cerevisiae* is known for its outstanding ability to produce ethanol in industry. Identifying the dynamic of gene expression in *S. cerevisiae* in response to fermentation is required for the establishment of any ethanol production improvement program. The goal of this study was to identify the discriminative genes between improved and repressed ethanol production as well as clarifying the molecular responses to this process through mining the transcriptomic data. Through 11 machine learning based algorithms from RapidMiner employed on available microarray datasets related to yeast fermentation performance under Mg^2+^ and Cu^2+^ supplementation, 172 probe sets were identified by at least 5 AWAs. Some have been identified as being involved in carbohydrate metabolism, oxidative phosphorylation, and ethanol fermentation. Principal component analysis (PCA) and heatmap clustering were also validated the top-ranked selective probe sets. According to decision tree models, 17 roots with 100% performance were identified. *OLI1* and *CYC3* were identified as the roots with the best performance, demonstrated by the most weighting algorithms and linked to top two significant enriched pathways including porphyrin biosynthesis and oxidative phosphorylation. *ADH5* and *PDA1* are also recognized as differential top-ranked genes that contribute to ethanol production. According to the regulatory clustering analysis, *Tup1* has a significant effect on the top-ranked target genes *CYC3* and *ADH5* genes. This study provides a basic understanding of the *S. cerevisiae* cell molecular mechanism and responses to two different medium conditions (Mg^2+^ and Cu^2+^) during the fermentation process.

## Introduction

In research and industry, *Saccharomyces cerevisiae* is used as one of the main microorganisms for bio-ethanol production. In addition to its high ethanol production capability, its stability for anaerobic fermentation and low pH tolerance facilitates its use in industry for ethanol production [1]. In terms of molecular biology, the genetics of *S. cerevisiae* is known, the genome has been sequenced, and many genes have been functionally annotated and characterized [2,3], so genetic manipulation of this organism is well developed [4]. There are different *S. cerevisiae* industrial strains used for bioethanol production. Molecular study of industrial strains with the aim of providing insight for improved ethanol production, is of great interest due to their importance for large-scale production. *S. cerevisiae JP1* is one of the dominant strains in fermentation industry since it exhibits high temperature tolerance, stability under low pH and high fermentation rate [5]. Several researches have been conducted on the *S. cerevisiae* metabolic engineering to generate efficient ethanol producing strains [6,7]. Suji et al [8], for example used the *PHO13* deletion in conjunction with *LAD1* and *ALX1* heterologous expression to improve *S. cerevisiae* for arabinose consumption, resulting in a 3.5-fold increase in specific ethanol productivity. Furthermore, transcriptomic studies have revealed the role of genes in ethanol fermentation. Under fermentation, gene expression analysis revealed the presence of stress-response and energy-related genes in *S. cerevisiae* supplemented with Mg^2+^ [9]. Upregulation of transketolase and transaldolase genes have been reported through transcriptome analysis of engineered *S. cerevisiae* under fermentation of arabinose sugar [10]. Identifying the molecular basis and dynamics of gene expression profiles related to yeast response in improved bioethanol production conditions is critical for developing new manipulated strains with increased ethanol yield. It also shed light on the mechanisms that yeast uses to improve production.

Metal supplements are effective in the yeast metabolic pathways that produce ethanol. Among these, zinc, magnesium, manganese, and copper have been extensively researched and shown to have regulatory effects on ethanol production [11, 12, 13]. Mg^2+^ ion is involved in phosphorylation, DNA and protein synthesis, as well as cell membrane rigidity and proliferation, and it has the potential to increase ethanol accumulation through fermentation [14, 9]. Furthermore, Mg^2+^ may improve the *S. cerevisiae* tolerance to high ethanol concentration during glucose and xylose fermentation [15, 16]. Mg^2+^ medium supplementation, in particular, resulted in a 29% increase in ethanol production by regulating the expression of cell wall and membrane related genes using *S. cerevisiae* [16]. Copper is also known as a critical element for yeast biological functions, particularly in its ion form Cu^++^. Some essential activities, such as cytochrome c oxidase, a component of oxidative phosphorylation, and superoxide dismutase are dependent on Cu^2+^ [17]. Copper stress, on the other hand, caused by an excess of copper, can result in ROS generation and DNA damage. At high concentrations, it also has a negative impact on cell membrane stability and enzyme activity. [18]. A high copper concentration (1.5 mM) inhibited cell growth, glucose and fructose consumption during fermentation by *S. cerevisiae* [19]. Few studies have been conducted to investigate the effect of Cu^2+^ on the physiology and fermentation ability of *S. cerevisiae* cell. As a result, despite their lack of research, Mg and Cu have the potential to modulate the gene expression network involved in the fermentation process.

It would be possible to identify the critical genes and clarify the mechanisms involved in the ethanol production process using bioinformatics-based analysis of the *S. cerevisiae* expression dataset. Computational approaches for identifying key genes involved in the fermentation process could elucidate the transcriptomic dynamics of yeast ethanol fermentation and reveal expression signatures that could be underutilized for improved production. RapidMiner is one of the most useful and widely used mining tools for data analysis [20]. Machine learning algorithms, both supervised and unsupervised models, are widely used in gene expression data analysis and gene identification [21,22]. Different gene selection algorithms, such as Information Gain, Information Gain Ratio, rule induction, SVM, and PCA, are widely used in gene expression analysis using RapidMiner. Cheng et al [23] used RapidMiner to preform four machine learning weighting models on gene expression datasets related to Huntington’s disease, including decision tree, rule induction, random forest, and generalized linear algorithms, in order to identify contributing genes to this disorder. In another study, Zinati et al [24] used ten different weighting algorithms to identify the genes that differentiate between sour and acidic lemon taste.

Valuable publicly available data on *S. cerevisiae* genome-wide expression experiments could be used for functional genomic analysis through machine learning. Machine learning algorithms’ discriminative ability aids in revealing the underlying biological process in microarray data analysis [20]. In light of the availability of such useful primary data sets and the potential of RapidMiner as an efficient tool for biological data analysis, we used available microarray expression dataset related to *S. cerevisiae* supplemented with Copper and Magnesium metal components under fermentation to investigate the underlying molecular basis of fermentation used by *S. cerevisiae*. The goal of this study was to identify the critical genes contribute to discriminate the improved (Mg^2+^ treatment) and low ethanol production (Cu^2+^ treatment at toxic concentration), as well as to elucidate the transcriptomic response of *S. cerevisiae* under these two conditions. *S. cerevisiae* transcriptome analysis using data mining and machine learning by both supervised and unsupervised models was used in this study as a novel approach to identify the underlying gene regulation mechanisms that can be used to optimize fermentation performance.

## Materials and Methods

### Data Collection

For this study available microarray datasets related to yeast fermentation performance under Mg^2+^ (500 mg/L) or Cu^2+^ (1 mg/L) supplementation was used. Microarray data of the industrial yeast *S. cerevisiae* JP1 strain downloaded from the GEO repository of the NCBI database (GEO number: GSE75803) was used. To meet the research objective, the probe sets with significant differential expression (concomitant Adj. p < 0.05 and B ≥ 3) were chosen for this study.

### Data cleaning

We used RapidMiner software (RapidMiner Studio 7.6) [25] to enter the 6300-differential expressed probe sets as numerical features, as well as high and low bioethanol as class features. For better processing, inefficient or redundant probe sets with less than or equal to a given standard deviation (SD) threshold (0.1), as well as correlated probe sets (correlation ≥ 0.95), were carefully removed from the dataset. The resulting list, which only contained efficient probe sets, was designated as the Final Cleaned (FCdb) database.

### Attribute weighting algorithms

Eleven attribute weighting algorithms with cut-off ≥ 0.7 were used in the FCdb to identify the most effective probe sets contributing to discriminate ethanol content. Weights close to 1 indicate that a specific probe set in ethanol content is more important. The main probe sets were those determined by the majority of AWAs (intersection of the weighing method). The attribute weighting algorithms used in this investigation, as well as the statistical background description for each one, are as follows (RapidMiner Studio 7.6):

### Weight by Information Gain and Information Gain Ratio

This algorithm is a well-established superior method for gene selection in microarray data analysis [26,27]. In this method, the attributes (probe sets) are weighted according to their class label (high or low ethanol production).

### Weight by Rule

Based on a single rule and the relationship between attributes (genes) and considering the errors, the weight of each attribute is measured through rule algorithm [28] and is used as a selective method for microarray analysis.

### Weight by Deviation, Weight by Correlation and Weight by Chi Squared Statistic

The standard deviation of attributes is used as a weighting parameter in the deviation weighting method. The correlation method, on the other hand, weighs the label attributes based on the correlation. In addition, for labeling the attributes, we used the Chi Squared Statistic weighting algorithm, which takes the Chi squared into account.

### Weight by Gini Index and Weight by Uncertainty

Due to the label attribute in this model, the weight of attributes is determined by measuring the Gini coefficient as an inequality index of sample data. According to each attribute, the lower the Gini index of the attribute, the more equal dispersion among attributes is considered. The weight for uncertainty model, on the other hand, is determined using the symmetrical uncertainty due to the class attribute.

### Weight by Relief

This model is one of the most reliable algorithms for weighting genes because it is based on the determination of values between probe sets of the same and different classes in a short distance.

### Weight by SVM

SVM is one of the most powerful classification models for gene expression analysis [21]. The SVM method weighs attributes using the coefficients of the normal vector of a linear SVM.

### Weight by PCA

This model performs attribute weighting due to the class attribute based on the component number parameter of PCA and the value of the components.

### Decision tree models

Eleven new datasets were generated using the entire probe sets with weight >0.70. They were annotated based on the models used for attribute weighting (Relief, Information gain, Uncertainty, Information gain ratio, Chi Squared, Rule, Correlation, Deviation, SVM and PCA, Gini index). Random Tree, Decision Tree, Decision Stump, and Random Forest were the tree induction models used for 12 datasets (FCdb and 11 datasets produced by specific weighting algorithms). Each model had four criteria (Gini Index, Gain Ratio Information Gain, and Accuracy). We used a ten-fold validation algorithm with appropriate sampling to create trees with RapidMiner. The performance of the model was evaluated and used to compare various models based on the accuracy of each model in identifying the target variable (high and low bioethanol content) and according to the attribute variables (normal expression of the probe set). Performance is expressed as a measure of model accuracy in this case. We calculated the accuracy by dividing the number of correct predictions by the total number of samples. The value of the attribute accuracy that is expected to be the same as the value of the labeled attribute is referred to as the correct prediction. These models were used with a minimum gain of 0.1 to obtain a split and a maximum tree depth of 20. For pruning 0.25 confidence level was considered with a pessimistic error calculation.

### Unsupervised analysis of the top ranked probe sets derived by supervised AWAs

Unsupervised principal component analysis (PCA) and hierarchical clustering heatmap were used to evaluate the power of top-ranked probe sets which differentiate the fermentation under different supplementation treatments. For unsupervised analysis, a web-based tool Clustvis (https://biit.cs.ut.ee/clustvis/) was used [30]. The PCA analysis was carried out in the PCA Methods R package using unit variance scaling on rows and Singular Value Decomposition (SVD) with the imputation method. The clustering heatmap was created with the pheatmap R package (version 0.7.7). The clustering heatmap was constructed using correlation, Pearson correlation subtracted from 1, and the average distance of all possible pairs [31].

### Kyoto Encyclopedia of Genes and Genomes (KEGG) pathway enrichment analysis

The pathway enrichment analysis was carried out using YeastEnrichr (https://maayanlab.cloud/YeastEnrichr/) [30,31]. The biochemical pathways related to key probe sets were identified using the KEGG2019 database. Pathways with p-value <0.1 were considered significant.

### Exploring for transcription factors among top-ranked genes and Regulator cluster analysis

We used yeastract database (http://www.yeastract.com/formrankbyhomotf.php) to identify transcription factors (TFs) among the 172 probe sets identified by at least 5 attribute weighting algorithms [32]. The TFs and their target genes were identified using this tool based on DNA binding sites and expression evidence. Furthermore, we used the regulator DB database (http://wyrickbioinfo2.smb.wsu.edu/cgi-bin/RegulatorDB/cgi/home.pl) to run regulator cluster diagram to determine the regulatory effect of the identified TFs on the target genes [33,34]. It provides data on mutant regulator expression for selected regulators and target genes.

## Results

### Ranking probe sets by AWAs

After cleaning 6300 probe sets by RapidMiner, we obtained 1813 probe sets. Eleven AWAs were used to identify informative probe sets. Following AWAs analysis, 172 probe sets were identified by at least 5 attribute weighting algorithms (Supplementary File, sheet S1). Furthermore, there were distinct probe sets classified by at least five algorithms that respond discriminatively to supplement treatment and/or are particularly related to ethanol production during fermentation. Sheet S2 of the Supplementary File contains the probe sets as well as the AWAs used to identify the probe sets. Some of the informative probe sets were recognized to be involved in carbohydrate metabolism, TCA cycle, oxidative phosphorylation, and ethanol fermentation while others were related to stress responses, cell membrane structure, and cell growth which could be indirectly effective in ethanol production. Some of the top informative genes are presented in (Table 1).

**Table 1.** Some of the informative probe sets identified by at least AWAs.

| probe sets | Standard Gene Name | AWAs Names | AWAs number | Gene Name |
| --- | --- | --- | --- | --- |
| A_06_P4554 | MRP8 | Chi Square Statistic, Correlation, Gini Index, Information Gain, Information Gain ratio, PCA, Relief, Rule, SVM, Uncertainty | 10 | Uncharacterized, response to stress |
| A_06_P1016 | OLI1 | Chi Square Statistic, Correlation, Gini Index, Information Gain, Information Gain ratio, PCA, Relief, Rule, SVM, Uncertainty | 10 | ATP synthase subunit 9, mitochondrial;OLI1;ortholog |
| A_06_P1397 | ADH5 | Chi Square Statistic, Correlation, Gini Index, Information Gain, Information Gain ratio, PCA, Rule, SVM, Uncertainty | 9 | Alcohol dehydrogenase 5;ADH5;ortholog |
| A_06_P1238 | PKC1 | Chi Square Statistic, Correlation, Gini Index, Information Gain, Information Gain ratio, PCA, Rule, SVM | 8 | Protein serine/threonine kinase |
| A_06_P3384 | GTR2 | Chi Square Statistic, Correlation, Gini Index, Information Gain, Information Gain ratio, PCA, Rule, SVM, Uncertainty | 9 | GTP-binding protein |
| A_06_P1063 | CYC3 | Chi Square Statistic, Correlation, Gini Index, Information Gain, Information Gain ratio, Relief, Rule, SVM, Uncertainty | 9 | Cytochrome c heme lyase |
| A_06_P1003 | COX1 | Chi Square Statistic, Correlation, Gini Index, Information Gain, Information Gain ratio, Rule, SVM, Uncertainty | 8 | cytochrome c oxidase |
| A_06_P2810 | PDA1 | Chi Square Statistic, Correlation, Gini Index, Information Gain, Information Gain ratio, Rule, SVM | 7 | Pyruvate dehydrogenase E1 component subunit alpha, mitochondrial;PDA1;ortholog |
| A_06_P2931 | QCR6 | Chi Square Statistic, Correlation, PCA, Rule, SVM | 5 | Cytochrome b-c1 complex subunit 6;QCR6;ortholog |
| A_06_P6820 | ALD6 | Chi Square Statistic, PCA, Rule, Uncertainty | 4 | Magnesium-activated aldehyde dehydrogenase, cytosolic;ALD6;ortholog |

### Decision Tree models

The decision tree models were used to achieve pattern recognition between important genes as well as with the genes with the highest distinguishing power. The lowest and highest performances were 0% and 100%, respectively (Supplementary File, sheet S3). There were 17 probe sets with 100% performance in the roots of decision tree models (Table 2).

**Table 2.** Decision Tree models roots identified as exhibited 100% performance.

| elements | GENE_SYMBOL | AWS | DESCRIPTION |
| --- | --- | --- | --- |
| A_06_P101<br>6 | OLI1 | 10 | BioProcess=ATP synthesis coupled proton transport |
| A_06_P247<br>5 | CWC21 | 9 | BioProcess=biological_process unknown |
| A_06_P100<br>2 | ORF:Q0017 | 9 | BioProcess=biological_process unknown |
| A_06_P103<br>4 | CYS3 | 9 | BioProcess=sulfur amino acid metabolism* |
| A_06_P113<br>1 | HTB2 | 8 | BioProcess=chromatin assembly/disassembly |
| A_06_P106<br>8 | KRE23 | 9 | BioProcess=biological_process unknown |
| A_06_P298<br>4 | CGR1 | 9 | BioProcess=rRNA processing* |
| A_06_P129<br>8 | ORF:YBR051W | 9 | BioProcess=biological_process unknown |
| A_06_P152<br>4 | ORF:YBR270C | 8 | BioProcess=biological_process unknown |
| A_06_P328<br>7 | ORF:YGR067C | 9 | BioProcess=biological_process unknown |
| A_06_P106<br>3 | CYC3 | 9 | BioProcess=not yet annotated |
| A_06_P205<br>1 | RPS13 | 9 | BioProcess=protein biosynthesis |
| A_06_P196<br>7 | OST4 | 10 | BioProcess=not yet annotated |
| A_06_P104<br>9 | ORF:YAL027W | 9 | BioProcess=biological_process unknown |
| A_06_P100<br>3 | COX1 | 8 | BioProcess=aerobic respiration |
| A_06_P202<br>3 | ORF:YDR036C | 7 | BioProcess=biological_process unknown |
| A_06_P102<br>3 | ORF:Q0297 | 8 | BioProcess=biological_process unknown |

### Unsupervised Analysis

As a complementary confirmation, the 172 top ranked probe sets were validated using PCA and hierarchical clustering heatmap, identified with supervised attribute weighting models. According to the results, the 172 significant probe sets could accurately differentiate between two different fermentation conditions, thus confirming the significance and accuracy of the identified probe sets (Fig. 1). In particular, the captured variances with the first two components on all recognized 6031 probe sets and informative 172 probe sets were up to 74% and 50%, respectively. Furthermore, it could efficiently separate informative 172 probe sets under Cu^2+^ or Mg^2+^ supplementation in the hierarchical clustering heat map, (Fig. 1). In total, 64 probe sets were up and down regulated by Mg^2+^ and Cu^++^, while 108 probe sets were up and down regulated through Cu and Mg supplementation, respectively (Fig. 2).

**Fig. 1.**
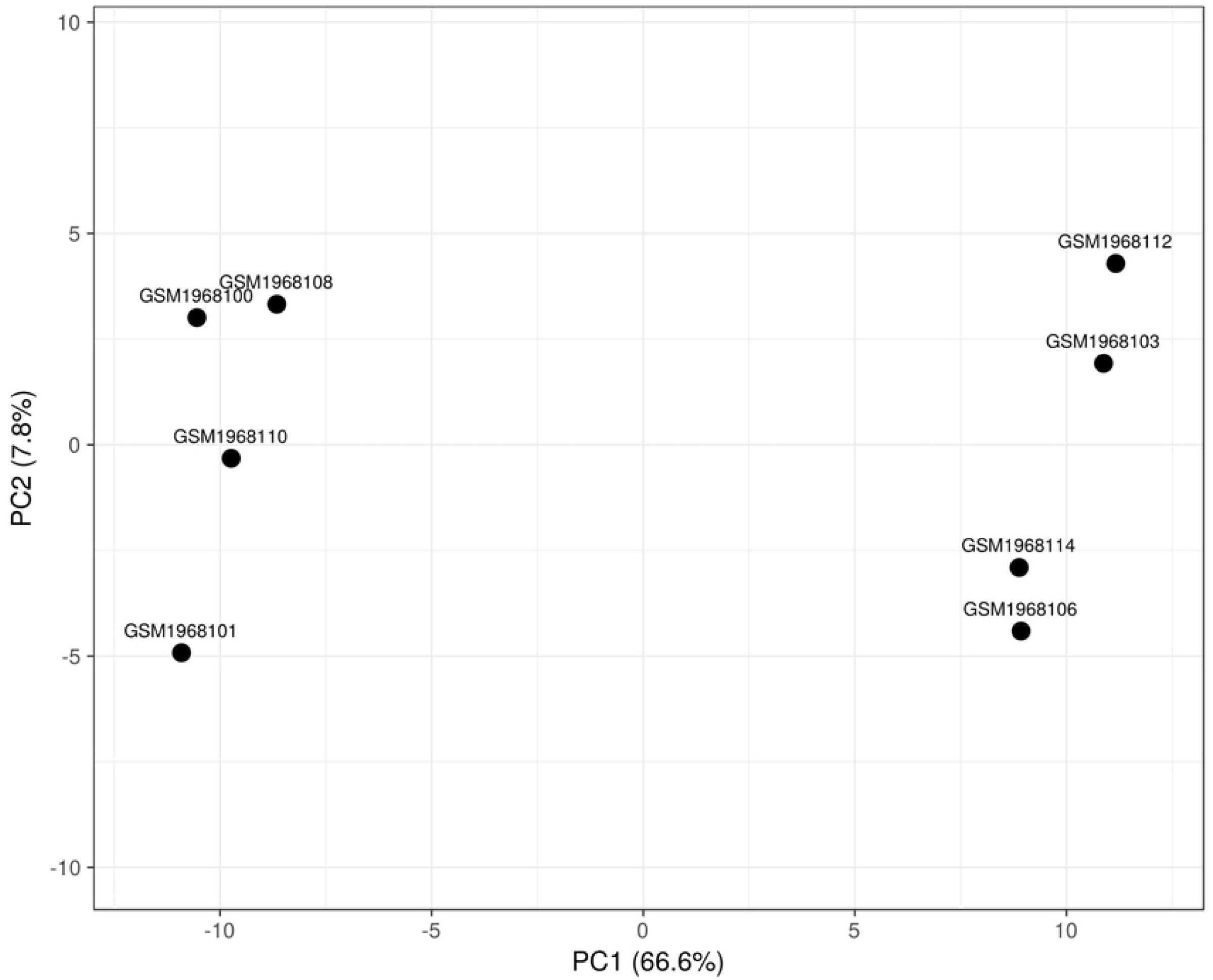
Two-dimensional plot related to the first two principal components. GSM1968101, GSM1968110, GSM1968100 and GSM1968108 are samples related to Mg^2+^ supplementation. GSM1968106, GSM1968114, GSM1968103 and GSM1968112 are samples related to Cu^2+^ supplementation.

**Fig. 2.**
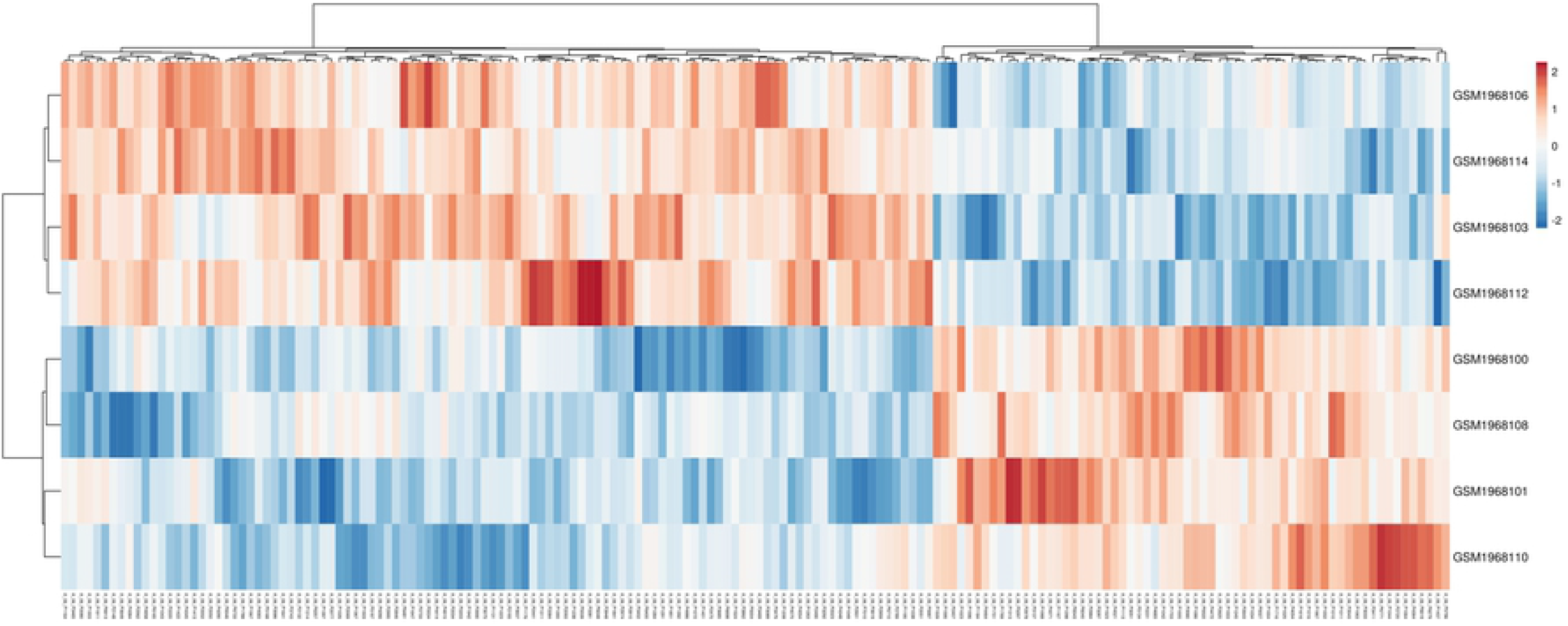
The heatmap related to 172 probe sets which were recognized by at least 5 attribute weighting algorithms (AWAs). Each row corresponds to the different samples including Mg^2+^ (high ethanol production) and Cu^2+^ supplementation (repressed ethanol production). Columns exhibits hierarchically clustered probe sets. The normalized intensity expressions of probe sets were shown as a color scale. The up and down-expression levels were represented as red and blue scales, respectively.

### Pathway enrichment analysis of genes

Significant enriched pathways such as Porphyrin metabolism, Oxidative phosphorylation, Glycolysis, Amino sugar and nucleotide sugar metabolism, Cell cycle, Meiosis and Citrate cycle (TCA cycle) were identified using the KEGG enrichment analysis. The enriched pathways and the related genes are presented in Table 3.

**Table 3.** KEGG enrichment analysis of 172 probe sets. The significant pathways with adjusted P-value < 0.1 are represented.

| Term | Adjusted P-value | Genes |
| --- | --- | --- |
| Porphyrin and chlorophyll metabolism | 0.000582985 | HEM2;HEM12;CYC3;YFH1 |
| Oxidative phosphorylation | 0.007415084 | OLI1;QCR6;ATP6;COX1;ATP2 |
| Endocytosis | 0.007415084 | CAP1;APL3;LAS17;ARC15;VPS25 |
| RNA degradation | 0.025376563 | POP2;RRP42;SSQ1;CCR4 |
| Meiosis | 0.049279656 | CLN3;HMRA2;MSN4;APC9;TPD3 |
| Autophagy | 0.049279656 | KCS1;VPS8;MSN4;PEP4 |
| Ubiquitin mediated proteolysis | 0.049279656 | UBC13;UBC6;APC9 |
| Protein processing in endoplasmic reticulum | 0.049279656 | OST4;UBC6;PDI1;SSE2 |
| Glycolysis / Gluconeogenesis | 0.064176371 | PDA1;PGM2;ADH5 |
| Galactose metabolism | 0.072088565 | GAL7;PGM2 |
| Phosphatidylinositol signaling system | 0.072088565 | KCS1;PKC1 |
| Amino sugar and nucleotide sugar metabolism | 0.075323668 | GAL7;PGM2 |
| Spliceosome | 0.075323668 | PRP43;ECM2;PRP8 |
| MAPK signaling pathway | 0.072088565 | TUP1;MKC7;MSN4;PKC1 |
| Pentose phosphate pathway | 0.075323668 | SOL4;PGM2 |
| Alanine, aspartate and glutamate metabolism | 0.075323668 | GDH3;NIT3 |
| Cell cycle | 0.075323668 | CLN3;TUP1;APC9;TPD3 |
| Citrate cycle (TCA cycle) | 0.075323668 | PDA1;LSC2 |
| Ribosome biogenesis in eukaryotes | 0.075323668 | UTP15;CKB1;RIO1 |
| Glycine, serine and threonine metabolism | 0.075323668 | SER1;CYS3 |

### Identification of transcription factors and their targets

Among the 172 informative probe sets identified by yeastract analysis were seven transcription factors: *YGR067C, HAP4, NRG2, TUP1, TOS8, MSN4, and PDC2*. Surprisingly, the targets of the identified transcription factors were discovered among the 172 genes identified by RapidMiner analysis and ranked by at least 5 algorithms (Supplementary File, sheet S5). These findings support the AWAs’ ability to correctly identify top-ranked probe sets. Furthermore, regulator clustering related to TFs and their targets (both ranked by at least eight algorithms) was performed to demonstrate the effect of top-ranked TFs on top-ranked target genes based on the transcription factors mutants. The results showed that *Hap4p, Tup1*, and *TOS8* mutants resulted in different ratios of up and down-regulation of target genes (Fig. 3). Although *Hap4* and *TOS8* resulted in down or up regulation of target genes, their effect on none of target genes was significant. According to the findings, the Tup1 transcription factor has the greatest impact on the target genes expression. The Tup1 knocked out mutant significantly induce the expression of *CYC3* (*YAL039C*), while causing highest level of down regulation of *YBL11C, ADH5* (*YBR145W*).

**Fig. 3.**
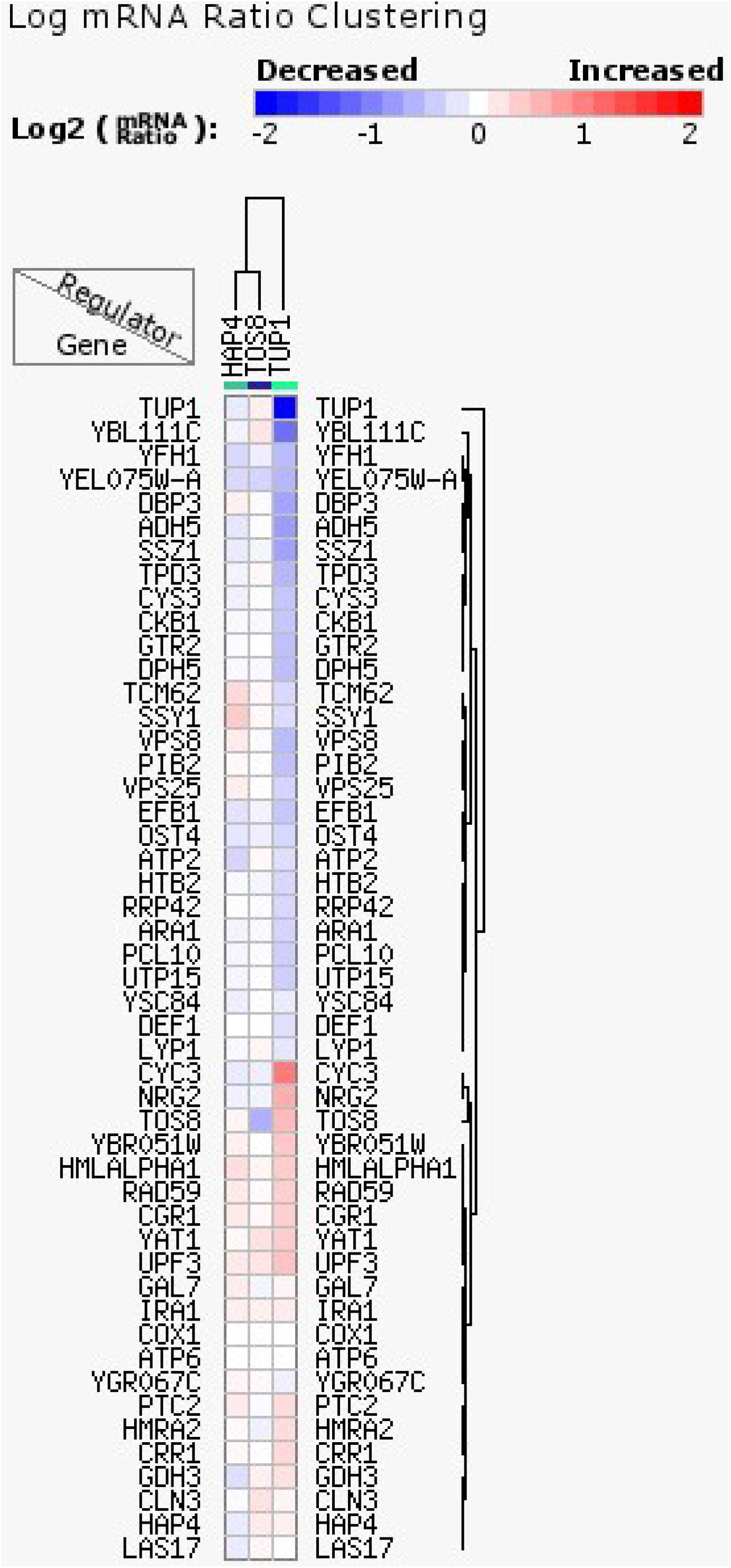
The regulatory clustering heatmap related to genes targeted by identified transcription factors *Hap4p* and *Tup1* and *TOS8*. The cluster is represented as the log mRNA ratio of each target gene in each regulator mutant.

## Discussion

In this study, machine learning and decision tree models were used to analyze the transcriptome of *S. cerevisiae* during the fermentation process in two conditions: repressed ethanol production and high ethanol production supplemented with Cu^2+^ and Mg^2+^. Indeed, for the most accurate prediction methods, we used both supervised and unsupervised models. In summary, we used 11 supervised models to achieve high accuracy results. In addition, a PCA analysis as an unsupervised model and a hierarchical clustering heat map were used to validate the 172 top-ranked probe sets identified by supervised-based models. Furthermore, we used pathway enrichment, transcription factor and regulatory analysis to validate the machine learning analysis results (Fig. 4). According to RapidMiner-assisted analysis, some probe sets were identified as playing a distinct role in ethanol production. Nonetheless, it should be noted that the function of some identified probe sets has not yet been clarified, despite the fact that they may be critical in ethanol production. *ADH5* or Alcohol dehydrogenase, which was weighted by 9 algorithms and classified in Glycolysis / Gluconeogenesis by KEGG enrichment analysis, contributes to ethanol production by reducing acetaldehyde to ethanol [35]. *OLI1* is distinguished by ten algorithms and is rich in Oxidative phosphorylation term, which encodes F0-ATP synthase subunit c and generates ATP in yeast mitochondria [36]. Metal ions, such as Cu^++^, are known to have a negative effect on mitochondrial respiratory components, as it slowed the respiratory chain in *PC12* and liver cells at toxic doses [37,38]. That is most likely the main reason for the down regulation of *OLI1*, which is an important component of the oxidative phosphorylation pathway when exposed to toxic Cu. RNA-seq analysis revealed that this gene was enriched as a significant gene between the wild and high glucose tolerant mutant strains of *S. cerevisiae* [39]. In addition to this gene, *COX1* has an AWA weight of 8 and is involved in the final electron chain reaction in the respiratory system [40]. It encodes one of the cytochrome c oxidase subunits and, like OLI1, has been shown to be repressed by Cu^2+^ treatment. *PDAI* encodes alpha subunit of pyruvate dehydrogenase and converts the pyruvate to acetyl-CoA through oxidative decarboxylation [41]. This gene was found to be enriched in the Glycolysis/Gluconeogenesis pathway by seven weighting algorithms used in this study. *PDAI* directs the pyruvate metabolism to Acetyl-COA in mitochondria to provide the TCA cycle substrate. In other words, directing the pyruvate to TCA cycle *PDAI* keeps pyruvate from being consumed in the fermentation process or ethanol production. *PDAI* was down regulated in Mg-containing medium, which accounts for improved ethanol production, and was upregulated in the repressed fermentation condition, by Cu. *QCR6* is a subunit of cytochrome bc1 complex and contributes to oxidative phosphorylation. Cytochrome C is known to be activated by Cu metal ion [42]. *QCR6* was up regulated, as expected, by Cu supplementation. Similarly, in Pichia stipites, cytochrome bc1 disruption resulted in increased ethanol production. [43]. Granados-Arvizu et al [44] also concluded that cytochrome bc_1_ complex repression would be a promising way to enhance ethanol production in *Saccharomyces stipitis. ALD6* or Aldehyde dehydrogenases is activated by Mg and have a distinct role in the formation of acetate from pyruvate in an alternate pyruvate dehydrogenase bypass pathway [45]. ALD6 expression was found to be increased with Mg supplementation, which corresponded to the activation of this enzyme by Mg++. Since it consumes the acetaldehyde source that ADH enzymes can use to produce ethanol, deleting *ALD6* via Crisper/CAS 9 genome editing resulted in increased ethanol production [46].

**Fig. 4.**
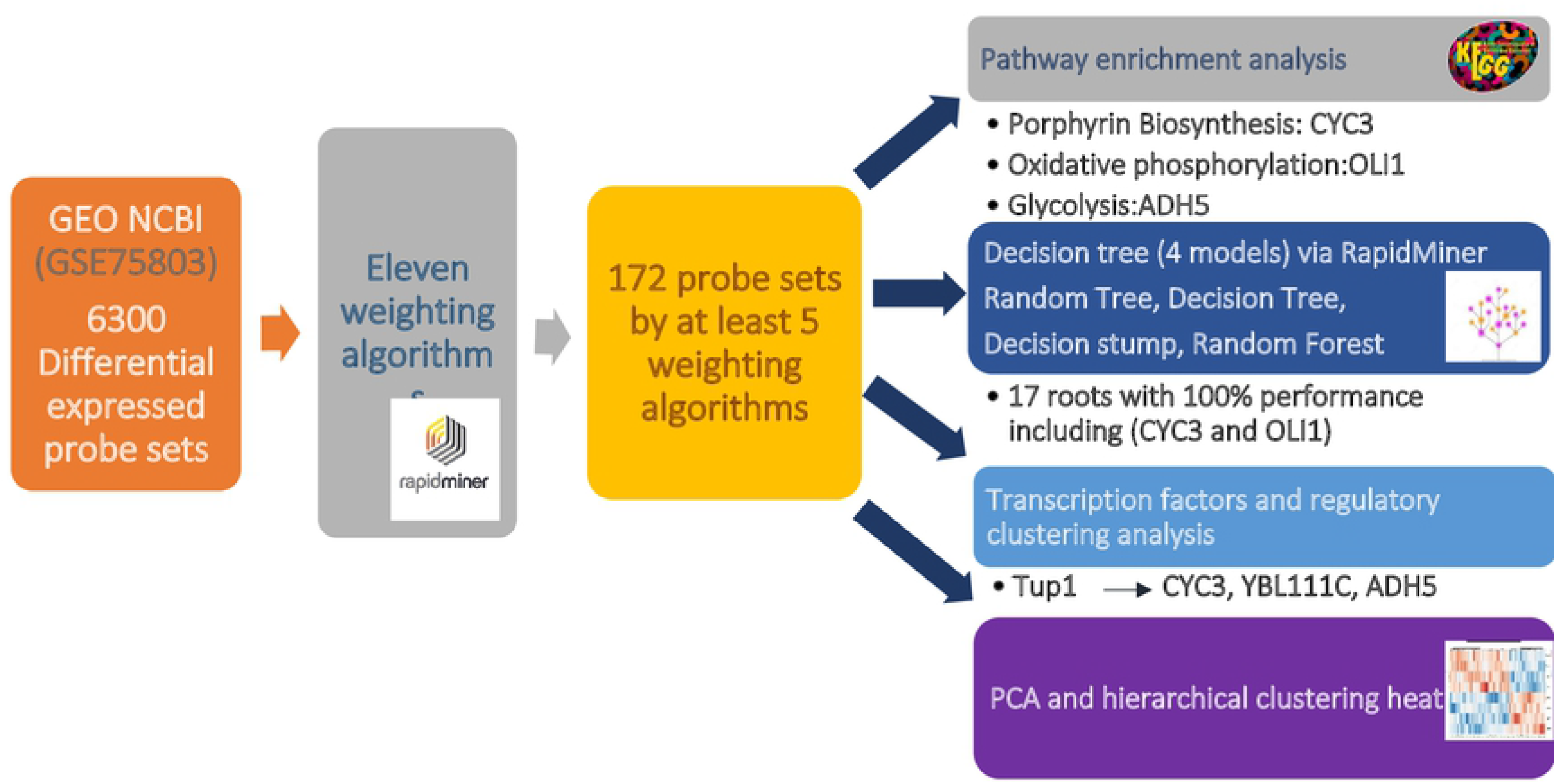
The schematic illustrates the methodology of the study with summarized results.

Based on decision tree analysis, 17 identified roots performed flawlessly, some of which have unknown molecular functions and have yet to be characterized. Surprisingly, *OLI1* and *CYC3* were identified by the highest attribute weighting algorithms (10 and 9), were enriched in the second most important biochemical pathway, and were also identified as decision tree model roots with 100% performance. *COX1* is also shown as a complete root, but it is identified using 8 weighting algorithms. As previously stated, *OLI1* is an F0-ATP synthase subunit c that contributes to the electron transport chain. *CYC3* is also known as Cytochrome c heme lyase and has a strong sensitivity to ethanol. Indeed, the null mutants for this gene showed ethanol sensitivity. Both of these genes are involved in ATP generation and are up regulated in Mg supplemented medium. Nonetheless, it has been established that Mg^2+^ has an effect on energy metabolism and ATP production in the cell [47].

Confirming the results of the AWAs and decision tree models analysis through RapidMiner, the Kegg pathway analysis showed that two most significant terms, porphyrin biosynthesis and oxidative phosphorylation, were enriched in *CYC3* and *OLI1* and *COX1*. Cell cycle and division, as well as Ribosome biogenesis, are identified as significant terms in the KEGG pathway enrichment. They may have an impact on ethanol production even though they do not directly contribute to the fermentation bioprocess. For example, in addition to its role in yeast cell growth and proliferation, which affects ethanol production, ribosome biogenesis is predicted to be associated with fermentation, and some related genes, such as *SFP1* are thought to be involved in glycolysis control as well [35,48]. Nonetheless, significant phosphatidylinositol signaling and MAPK signaling pathways identified in this study by enrichment analysis were reported to be responsible for cell proliferation/growth regulation and critical for stress responses [49,50]. *PKC1* which was attributed by 8 algorithms and remarkably enriched in phosphatidylinositol signaling system is a serine/threonine kinase which is suggested to have role in response to copper toxicity since it was upregulated in Cu^2+^ supplementation or reduced ethanol production according to heatmap clustering. Confirming this finding, Zhou et al [49] reported that 5-hydroxymethyl-2-furaldehyde, which is toxic to industrial fermentative *S. cerevisiae* strain, increases the expression of *PKC1* gene. Furthermore, according to AWAs analysis, some genes are involved in stress responses, cell growth and proliferation, protein synthesis, fatty acids and lipid metabolism, all of which may contribute to ethanol production efficiency. *MRP8* was assigned by ten algorithms as a response to cell wall stress, and its expression has been reported to be induced under stress conditions [51]. Its function, however, is unknown. Cu supplementation induces the expression of this gene in response to the stress condition caused by copper. *GTR2*, a GTPase subunit, was weighted using nine algorithms. It is suggested in this study that it contributes to tolerance response to CU inhibitor because it was up regulated by copper. As an implication for this result, the null mutant related to *GTR* gene showed decreased resistance to Zn metal at inhibitory amount [52].

According to the crucial role of TFs in gene expression regulation and to confirm the results obtained from attribute weighting algorithms analysis, the TFs and their targets were explored among 172 probe sets. According to the regulatory clustering analysis, *Tup1* has a significant effect on the top-ranked target genes. *Tup1* is a transcriptional repressor in *S. cerevisiae* has the ability to repress target genes via various molecular mechanisms, and it contributes to carbon catabolite repression of transcription by glucose [53, 54]. Regarding the results of this study on regulatory clustering analysis, the *Tup1* mutant caused decreased expression in some of the target genes and up regulation in others. In other words, the deletion of *Tup1* resulted in downregulation of *YBL111C*, whose biological function is unknown and *YBR145W (ADH5)* at most. The *ADH5* gene has also been identified as the top-ranked gene with 9 AWAs through RapidMiner analysis. On the other hand, the *TUP1* knock out resulted in significant upregulation of *YAL039C* (*CYC3*). Indeed, the *CYC3* gene, which was confirmed by the greatest number of AWAs and a decision tree model, was also shown to be a top target of the transcription factor involved in ethanol production responses in this study. *Hap4* is a transcription factor involved in the regulation of the respiratory genes’ expression and ethanol tolerance. The role of *TUP1* and *HAP4* in glucose fermentation have been studied and recently confirmed in thermotolerant yeast, *Ogataea polymorpha* [54]. Moreover, the overexpression of *HAP4* gene caused enhanced glucose consumption and ethanol production in *S. cerevisiae* [55,56]. In this study, the HAP4 gene was also identified as top-ranked gene attributed by nine AWAs. Although the results confirm its involvement in the identified probe sets regulation, it does not demonstrate significant up or down-regulation effect on the target genes. Overall, *OLI1, CYC3, COX1 and ADH5* were ranked as the most critical genes in the differentiation of two improved and repressed ethanol production conditions because they were the most frequently identified genes across analyses. These important findings shed light on the complex pathways and regulatory responses that genes use to contribute to ethanol production. However, additional experimental analysis could fully clarify the results. Overall, the findings of this study could be used to further investigate the possibility of improving ethanol through overexpression or knock out strategies. Furthermore, additional experimental testing to confirm the findings is strongly advised.

## Abbreviations

(PCA): Principal component analysis
(KEGG): Kyoto Encyclopedia of Genes and Genomes
*ADH5*: (Alcohol dehydrogenase)
(CYC3): Cytochrome c heme lyase

## Declarations

### Ethics approval and consent to participate

Not applicable

### Consent for publication

Not applicable

### Availability of data and materials

All data generated or analyzed during this study are included in this published article [and its supplementary information files}.

### Competing interests

The authors declare that they have no competing interests

### Funding

No funding

### Authors’ contributions

SS, ZZ and AN contributed to the study conception and design. ZZ and SS analyzed the Data. The manuscript was written by SS. ME, ZZ and AN contribute to scientific revision of the manuscript. All authors read and approved the final manuscript.

## Acknowledgements

Not applicable

## Supplementary data

The Supplementary File, sheet S1 and S2. The list of the probe sets along with the attribute weighting algorithms (AWAs) through which the probe sets were identified.

The Supplementary File, sheet S3 and S4. The decision tree models (performance and roots).

The Supplementary File, sheet S5. The list of identified transcription factors and their targets.

## Notes

### Competing Interest Statement

The authors have declared no competing interest.

